# A CRISPR-based instant DNA repositioning system and the early intranuclear life of HSV-1

**DOI:** 10.1101/2022.04.08.487454

**Authors:** Zhaoyang Fan, Juan Xiang, Pei Xu

## Abstract

The intranuclear localization of viral DNA genomes in relation to the intranuclear environment plays critical roles in determining virus fate. Recent advances in the application of chromosome conformation capture-based sequencing analysis (3C technologies) have revealed valuable aspects of the spatiotemporal interplay of viral genomes with host chromosomes. However, to elucidate the causal relationship between the subnuclear localization of viral genomes and the pathogenic outcome of the infection, manipulative tools are needed. Instant repositioning of viral DNAs to specific subnuclear compartments amid infection is a powerful approach to synchronize and interrogate this dynamically changing process in space and time. Herein, we report an inducible CRISPR-based two-component platform that relocates extrachromosomal DNA pieces (5 kb to 170 kb) to the **nu**clear **p**eriphery **in**stantly (CRISPR-nuPin). Based on this system, investigations of herpes simplex virus 1 (HSV-1), a prototype member of the human herpesvirus family, revealed unprecedently reported insights into the early intranuclear life of the pathogen: I) Viral genomes tethered to the nuclear periphery upon entry, compared with those in the nuclear interior, were wrapped around histones with increased suppressive modifications and subjected to stronger transcriptional silencing and prominent inhibition. II) Relocating HSV-1 genomes at 1 hour post infection significantly promoted transcription of viral β and γ genes, termed an “Escaping” effect. III) Early accumulation of ICP0 was a sufficient but not necessary condition mediating “Escaping”. IV) Subnuclear localization was only critical during early infection. Importantly, the CRISPR-nuPin tactic should be widely applicable to many DNA viruses.

**Summary:** The intranuclear localization of viral DNA genomes plays a critical role in determining virus fate. To elucidate the causal relationship between subnuclear localization and the pathogenic outcome of DNA viruses, manipulative tools are needed. Herein, we report an inducible CRISPR-based two-component platform that relocates DNA pieces (5 kb to 170 kb) to the **nu**clear **p**eriphery **in**stantly (CRISPR-nuPin). Utilizing this tactic, we interrogated the early intranuclear life of herpes simplex virus 1 (HSV-1), a prototype of human herpesviruses, in space and time and revealed that I) viral genomes tethered to the nuclear edge upon entry were prone to suppressive histone packaging and severe inhibition. II) Relocating HSV-1 genomes to the nuclear fringe at 1 hour post infection promoted transcription of viral genes (“Escaping”). III) Early accumulation of ICP0 was a sufficient but not necessary condition mediating “Escaping”. IV) Subnuclear localization was a critical factor only during early infection.

**Highlights:**

- CRISPR-nuPin is an inducible two-component DNA repositioning system
- It mediates instant nuclear edging of viral DNA during infection
- A powerful approach to interrogate DNA viruses in space and time
- Viral DNA at the nuclear periphery upon entry is strongly silenced

**In brief:** An inducible two-component CRISPR-based platform that instantly repositions HSV-1 genomes to the nuclear edge unveils intranuclear space heterogeneity for the incoming viral genomes and dynamic stages of the host-virus interplay during early infection of the pathogen.

## Introduction

Cellular chromosomes are not randomly positioned in the nucleus, and the dynamic organization of intranuclear DNA in space and time plays a fundamental role in regulating the cellular transcriptome. Recent innovations in mapping and manipulative technologies have boosted scientific enthusiasm to address the cause-effect relationship between cellular gene transcription regulation and its relative intranuclear locus on the 4-dimensional scale [1–3]. Appreciation of the complexity and the critical transcriptional regulatory role of the higher-order organization of the host genome raises enormous interest in the field of virology, believing that DNA viruses, co-evolved parasites of the host machineries, are interrelated with and/or taking advantage of the cellular 3D genome organizations. Currently, evidence supporting this concept mostly manifests in descriptive experiments identifying and/or characterizing host genomic hot spots in association with viral genomes using immunofluorescence staining and fluorescent *in situ* hybridization (FISH) or, more recently, by chromosome conformation capture (3C)-based methods [4–8]. Few investigations addressing the causal relationship underlying intranuclear host-virus spatial correlations have been conducted, partially due to a lack of proper probing tools. In this study, we took advantage of the advent of advances in CRISPR-based genome re-organization tools and constructed an inducible two-component CRISPR-nuPin platform that enabled repositioning of extrachromosomal DNA pieces to the nuclear edge within minutes. Furthermore, we used the system to interrogate the early intranuclear life of HSV-1, a prototype medium-sized DNA virus, in space and time.

HSV-1, a member of subfamily *Alphaherpesvirinae*, family *Herpesviridae*, is a double stranded DNA virus of approximately 153 kb in size that infects over 66.7% of the population in 0-49-year-olds globally [9]. A mature, infectious virion of HSV-1 consists of a virus envelope, tegument layer and an icosahedral capsid encapsdating one copy of the viral genome. HSV-1 uses the strongest molecular motor known (portal vertex) to pack its negatively charged genomes into a relatively tiny space, resulting in tens of atmospheres of pressure inside the capsid [10–13]. The viral capsid docks itself to the nuclear pore complex (NPC), and the internal pressure provides an essential driving force for viral genome release and transportation through the NPC in a rod-like structure [10, 11, 14–17]. Upon genome ejection into the nucleus, HSV-1 adopts an active lytic replication cycle in almost every susceptible cell line *in vitro*, and three sets of viral genes are expressed in a tightly regulated chronological order, termed immediate early (α), early (β) and late genes (γ_1_, γ_2_). The immediate events postnuclear entry of viral genomes center around the raging competition between the urge of viral genes to express and the repressive forces from the host cell to silence these exogenous DNAs. Abundant evidence shows that nuclear HSV-1 genomes are immediately sensed, assembled with histones, associated with *de novo*-formed nuclear bodies (PML bodies), and loaded with host proteins, including repressor complexes, to restrict viral transcription and defend against infection [18–21]. To counteract host intrinsic immunity, HSV-1 viral tegument protein αTIF(VP16) and its associated complex are transported into the nucleus independently of the viral capsid and initiate efficient transcription of α genes, including infected cell protein 0 (ICP0) [22–26]. ICP0 is a multifunctional E3 ligase and promiscuous transactivator essential for the expression of post-α genes (β and γ genes) at a low multiplicity of infection [27]. The protein seemed to execute transcription-relevant functions in tandem: first, the dispersal of PML bodies through proteasome-dependent degradation of PML and Sp100 and then inactivation of the HDAC1-2-CoREST-REST-LSD1 repressor complex [18, 28, 29]. An important but blurred aspect of these elegantly elaborated molecular mechanisms is the precise sequential order of these events and their relationship with the diverse spatial localizations of the viral genomes in the host nuclei.

Herein, utilizing the CRISPR-nuPin platform, we took a very first step addressing the complicated and heterogeneous host-virus interactions on a 4-dimensional scale and revealed several intriguing aspects of the early intranuclear life of HSV-1. In summary, we report that I) the CRISPR-nuPin strategy is a two-component platform consisting of a dCas9-emerin fusion protein and sgRNAs. II) CRISPR-nuPin drove relocation of extrachromosomal DNA pieces up to 170 kb to the inner nuclear margin efficiently and fleetly upon electroporation of sgRNAs. CRISPR-nuPin can reposition prey DNA via a single sgRNA targeting site and thus has wide application to various DNA viruses replicating in the nucleus. III) HSV-1 genomes deposited to the nuclear edge immediately upon entry were subjected to stringent transcriptional silencing and major growth inhibition, up to a multiplicity of infection (MOI) of 20. This subnuclear localization bias was only detected during the immediate early period of infection. IV) Dislodging HSV-1 genomes from their originally occupied spatial niches to the nuclear margin at 1 hpi significantly promoted viral gene expression and virus production (“Escaping”). V) Early accumulation of ICP0 protein was a sufficient but not necessary condition in “Escaping”. VI) Shortly post infection, HSV-1 genomes are no longer sensitive to subnuclear localization changes.

## Material and Methods

### Plasmids and Transfection

The emetine coding sequence was amplified from the cDNA of HEp-2 cells. dCas9 (D10A/N863A) was amplified from the lentiSAMv2 backbone (Addgene, #75112). The coding sequences of dCas9 and emerin were inserted into a pcDNA3.1 vector and arranged as NLS (nuclear localization signal)-dCas9-NLS-emerin-Flag-T2A-GFP under a CMV promoter for transient transfection. The NLS-dCas9-NLS-emerin-Flag-T2A-GFP was cloned into the all-in-one doxycycline inducible lentiviral vector pCW57-MCS1-2A-MCS2, a gift from Pro. Deyin Guo’s lab (Sun Yat-sen University) for lentivirus packaging and transduction of HEp-2 cells. The 170 kb plasmid used in Figures 1 and 2 was a bacterial artificial chromosome (BAC) containing the HSV-1 (F strain) genomic sequence (BAC-HSV-1). The 5 kb plasmid used in Figures 1 and 2 was a modified pcDNA3.1 vector. All constructs were verified by sequencing.

**Figure 1.**
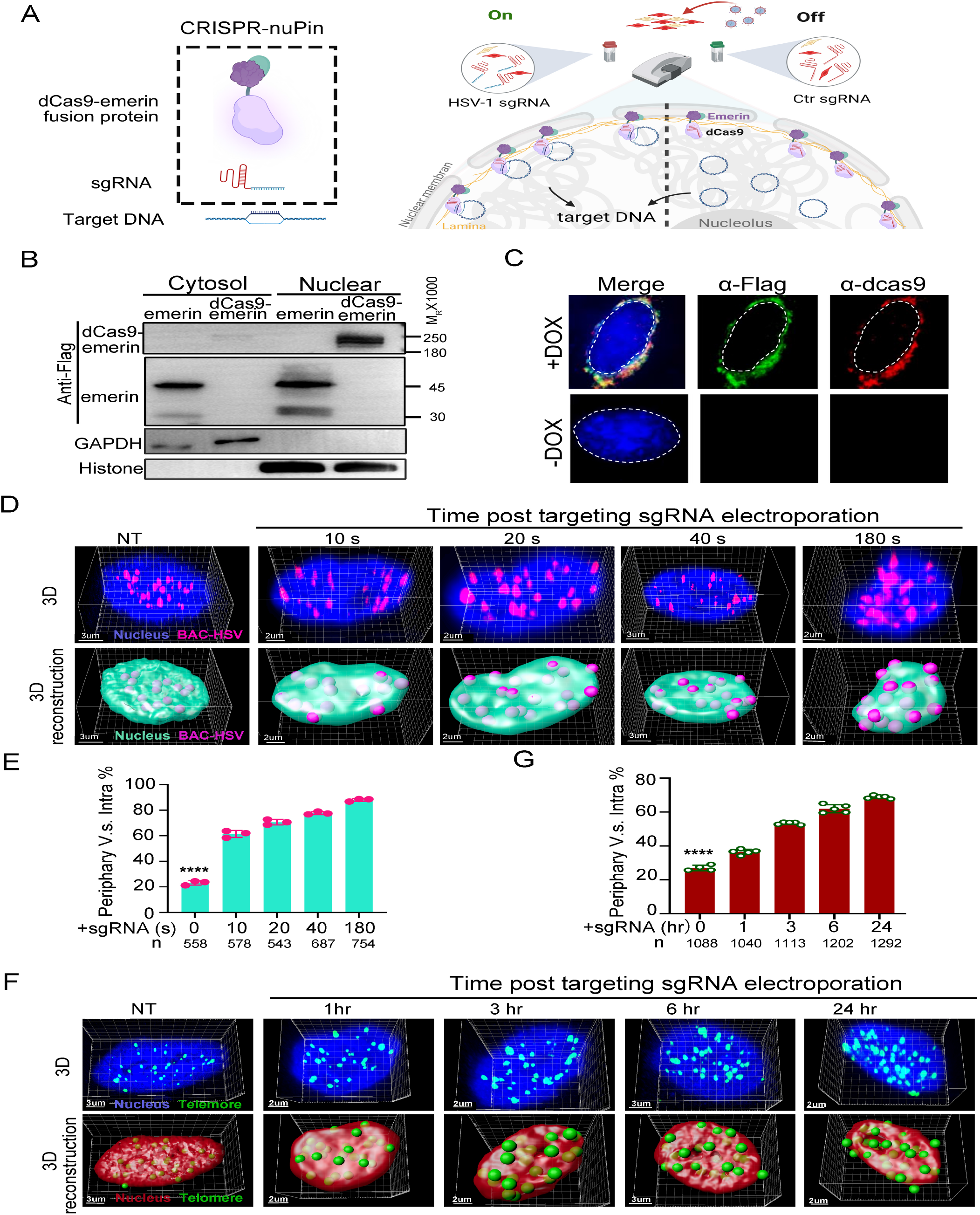
Validation of the two-component CRISPR-nuPin system. **A.** Schematics of the CRISPR-nuPin system. **B.** Subcellular localization of NLS-dCas9-NLS-emerin-Flag-T2A-GFP and emerin-Flag in transiently transfected HEK293T cells. Immunoblotting was performed using anti-Flag, anti-GAPDH and anti-histone antibodies. **C.** dCas9-emerin cells were mock treated (-DOX) or treated with doxycycline (+DOX), and the intracellular distribution of dCas9-emerin protein was detected by immunofluorescence staining with anti-Flag (green) and anti-dCas9 (red) antibodies. dCas9-emerin cells (+DOX) containing 170 kb BAC plasmids were electroporated with BAC targeting sgRNA (**D, E**), or dCas9-emerin cells (+DOX) were electroporated with telomere targeting sgRNA (**F, G**). At the indicated time points post electroporation, the cells were fixed, and the BACs or telomeres were stained by FISH and scanned under a Nikon microscope (N-STORM). Representative Z series images are shown in the upper panel of D and F, and 3D reconstruction of the images is displayed in the bottom panel of D and F. The percentage of BAC stains (**E**) or telomere stains (**G**) located at the nuclear edge versus total intranuclear stains at each time point was calculated and plotted. Total counted stains at each time point are shown as n in the bottom of E and G.

**Figure 2.**
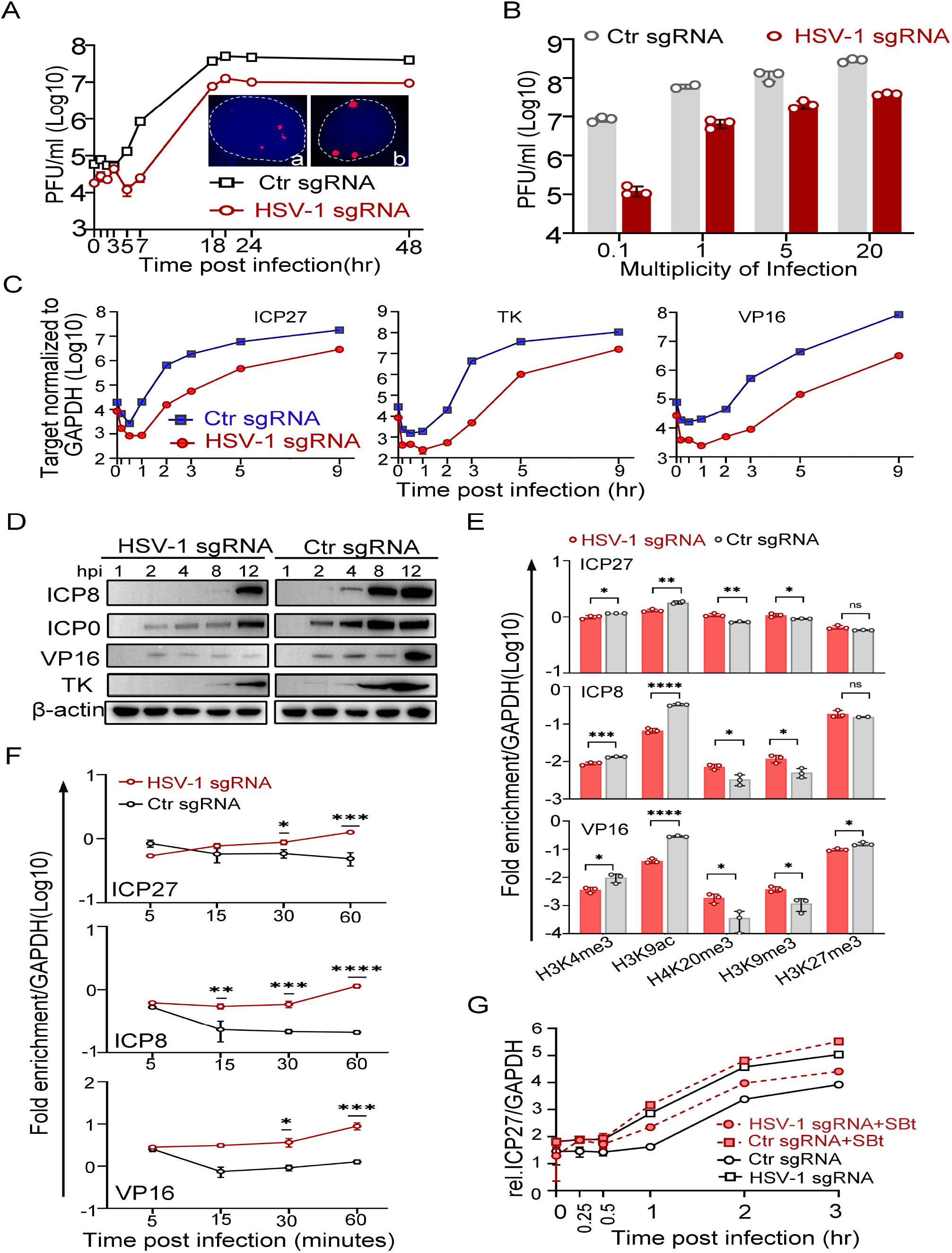
Transcription from HSV-1 genomes deposited to the nuclear edge upon entry was strongly inhibited. **A-F.** dCas9-emerin cells transfected with plasmids expressing HSV-1 sgRNA or Ctr sgRNA for 24 hr were infected with HSV-1 at an MOI of 1 (A), the indicated MOIs (B) or an MOI of 5 (C to F). **A.** Single-cycle growth kinetics of HSV-1 by plaque assay. **B.** Cell-associated HSV-1 virus was titrated by plaque assays at 12 hpi. **C.** Total RNA was extracted at 0, 0.17, and 0.5 1, 2, 3, 5 and 9 hpi, and the mRNA levels of ICP27, TK, ICP0 and VP16 were measured by qPCR (0.17 and 0.5 hpi were not marked). **D.** The viral protein levels of ICP0, ICP8, TK, and VP16 at the indicated time points post infection were examined by immunoblotting. β-actin served as a loading control. **E.** ChIP assay of differentially modified histones on the promoters of ICP27, ICP8 and VP16 of HSV-1 in HSV-1 sgRNA- or Ctr sgRNA-expressing dCas9-emerin cells infected with HSV-1 at 0.5 hpi performed using antibodies specific for H3K4me3, H3K9ac, H3K9me3, H4K20me3 and H3K27me3. **F.** ChIP assay of heterochromatins on the promoters of ICP27, ICP8 and VP16 of HSV-1 in HSV-1 sgRNA- or Ctr sgRNA-expressing dCas9-emerin cells infected with HSV-1 at the indicated time points was performed using an antibody against H3K9me3. For E and F, the results were expressed as enrichment (fold) by comparing the fraction of viral promoters immunoprecipitated by the indicated antibodies to the fraction of GAPDH immunoprecipitated in the same reaction. **G.** HSV-1 sgRNA- or Ctr sgRNA-expressing dCas9-emerin cells were treated with SBt for 5 hr and infected with HSV-1 at an MOI of 1. The mRNA level of ICP27 at the indicated time points was measured by qPCR.

Transient transfection of HEK293T cells used standard polyethylenimine (PEI) (Sigma, #408727). For transfection of HEp-2 cells and other mammalian cell lines, Jetprime (Polyplus, #101000046) was used according to the manufacturer’s protocol.

### Cells, viruses, and infection

HEp-2, HEK293T and Vero cells were grown in Dulbecco’s modified Eagle’s medium (DMEM, Corning, #10013074) supplemented with 10% fetal bovine serum (FBS, Gibco, #42G4086K). U2OS cells were cultured in McCoy’s 5A (Procell, #PM150710) with 10% FBS. dCas9-emerin cells were established by lentivirus transduction of HEp-2 cells, selected under puromycin (1 μg/ml) (Gibco, #A1113803), induced by doxycycline (DOX) at 3 μg/ml and sorted for high GFP-expressing cells by MoFLO Astrios EQs (Beckman Coulter Life Sciences) using 488 nm lasers. dCas9-emerin cells were treated with DOX for at least 3 days before HSV-1 infection experiments.

HSV-1(F) and recombinant HSV-1 carrying GFP gene (R8515) virus were prepared in HEp-2 cells and titrated in Vero cells by plaque assay. Recombinant HSV-1 lacking the ICP0 gene (ΔICP0) was amplified and titrated in U2OS cells by plaque assay. Viruses were stored at −80°C as single-use aliquots.

Except for the plaque assay and virus amplification, all infections were conducted by inoculating cells with HSV-1 virus at the desired MOI on ice for 2 hr (referred to as 0 hpi). Infected cells were cultured in DMEM supplemented with 1% FBS at 37°C. At the indicated time points, the HSV-1 titer was determined by collecting all infected cells, freeze and thawing three times, sonicating them at 20% amp for 7 s and titrating them into Vero cells.

### Lentivirus production

To produce lentivirus, HEK293T cells were seeded in 60 mm dishes and transfected with packaging plasmids pCMV-dR8.91 and VSV-G and a lentiviral backbone plasmid containing the dCas9-emerin-flag-T2A-GFP insertion or GFP insertion only at an appropriate ratio. The medium was changed at 6-8 hr post transfection. Culturing medium was collected at 48 hr post transfection, filtered through 0.45 μm filters and centrifuged at 1000 g for 10 min to remove cell debris. The supernatant containing lentiviruses was added freshly to cells for infection or frozen at −80 °C.

### Fluorescence in situ hybridization (FISH)

HSV-1 DNA FISH probes were prepared from BAC-HSV-1 through nick translation (exon biotechnology, #21076) with Cy3-labeled dNTPs. Then, 5 kb plasmid DNA FISH probes were prepared similarly from the plasmid through nick translation. Briefly, 2 μg of template DNA was mixed with Cy3-labeled dNTPs, nick translation buffer, nuclease-free water, and nick translation enzyme according to the manufacturer’s protocol. Then, the mixture was incubated for 1-5 hr at 15°C. The labeled probe was precipitated and redissolved in nuclease-free water. The Telomere FISH probe was purchased from PNA (TelG-Alexa488, #F1008). The hybridization procedure was slightly modified from a previously described protocol [30]. In brief, the cells were fixed with 4% paraformaldehyde (Sigma, #P6148) at room temperature (RT) for 15 min, incubated with 0.1% Tris-HCl pH 7.0 for 10 min, prepermeabilized with 0.8% Triton (Sigma, #T8787) in PBS at RT for 10 min, incubated with 20% glycerol (Sangon Biotech, #A600232-0500) in PBS for 20 min, permeabilized with 0.8% Triton PBS for 30 min, treated with RNase A (Omega, #L15UM) at 37°C for 50 min, and incubated with 50% formamide (VETEC, #V900064) in 2X saline-sodium citrate (SSC) buffer (0.3 M NaCl, 30 mM sodium citrate) for 10 min, with PBS washes between steps. The cells were incubated with probes diluted to 20 ng/ml in hybridization solution (2XSSC, 50% formamide, 10% dextran sulfate (Sigma, #D8906), 1% Triton) at 85°C for 10 min and 37°C for 20 hr. After hybridization, the cells were sequentially washed with PBS containing 75%, 50% and 25% wash buffer (2XSSC, 70% formamide, 10% dextran sulfate).

For FISH and immunofluorescent staining, cells were further treated and stained as described in the immunofluorescent staining section.

### Quantitative Real-Time PCR

Total RNA was extracted using an E.Z.N. A Total RNA Kit I (OMEGA, #R6834-02), followed by cDNA synthesis using the Evo M-MLV RT Kit (Accurate biology, #AG11603). Total DNA was isolated using a Virus DNA Kit (OMEGA, #D3892-01). qPCR was performed using the Sybrgreen detection systems (Accurate biology, #AG11701) on the StepOne™ system (ThermoFisher). GAPDH was used as a loading control.

The primers used are as follows: ICP0 forward, 5’-ACTGCCTGCCCATCCTG GACA-3’; ICP0 reverse, 5’-CCATGTTTCCCGTCTGGTCC-3’; ICP27 forward, 5’-CGGGCCTGATCGAAATCCTA-3’; ICP27 reverse, 5’-GACACGACTCGAAC ACTCCT-3’; TK forward, 5’-CCAAAGAGGTGCGGGAGTTT-3’; TK reverse, 5’-CTTAACAGCGTCAACAGCGTGCCG-3’; VP16 forward, 5’-CCATTCCACCA CATCGCT-3’; VP16 reverse, 5’-GAGGATTTGTTTTCGGCGTT-3’; Us1 forward, 5’-ATCAGCTGTTTCGGGTCCTG-3’; Us1 reverse, 5’-TCGGCAGTATCCCA TCAGGT-3’; Us12 forward, 5’-AACGCACCAAACAGATGCAG-3’; Us12 reverse, 5’-CGTCCAAACCCACCGACATA-3’; human GAPDH forward, 5’-GAAGGT GAAGGTCGGAGTC-3’; and human GAPDH reverse, 5’-GAAGATGGTGATG GGATTTC-3’. ICP27 promoter forward, 5’-CCGCCGGCCTGGATGTGACG-3’; ICP27 promoter reverse, 5’-CGTGGTGGCCGGGGTGGTGCTC-3’; ICP8 pro moter forward, 5’-CCACGCCCACCGGCTGATGAC-3’; ICP8 promoter reverse, 5’-TGCTTACGGTCAGGTGCTCCG-3’; VP16 promoter forward, 5’-GCCGCC CCGTACCTCGTGAC-3’; VP16 promoter reverse, 5’-CAGCCCGCTCCGCTT CTCG-3’; GAPDH promoter forward, 5’-TTCGACAGTCAGCCGCATCTTCTT-3’; GAPDH promoter reverse, 5’-CAGGCGCCCAATACGACCAAATC-3’.

### Cell viability assay

Cell viability assays were performed using a Cell Counting Kit-8 (CCK-8, Bimake, #B34302) according to the manufacturer’s instructions. The fluorescence intensity was measured in a Synergy H1 microplate reader (Biotek) at 450 nm. Wells containing only culturing medium served as controls.

### Subcellular fractionation

Subcellular fractionation was performed as previously described [31]. In short, 1X10^6^ cells were collected, pelleted by centrifugation at 1000 g for 5 min, resuspended in 0.3 ml of ice-cold buffer 1 (150 mM NaCl, 50 mM HEPES [pH 7.4], 25 μg/ml digitonin, 10 μl/ml protease inhibitor), incubated for 30 min at 4°C and then centrifuged at 4,600 rpm for 5 min. The supernatant was collected, and the pellets were washed and resuspended in 0.3 ml of ice-cold buffer 2 (150 mM NaCl, 50 mM HEPES [pH 7.4], 1% [vol/vol] NP-40, 10 μl/ml protease inhibitor) and incubated for 30 min on ice. The samples were centrifuged at 8,700 rpm for 5 min to pellet nuclei, and the supernatants were collected and combined with the previous collection, representing the cytosol. The pellets were washed and resuspended in 0.2 ml of ice-cold buffer 3 (150 mM NaCl, 50 mM HEPES [pH 7.4], 0.5% [wt/vol] sodium deoxycholate, 0.5% [wt/vol] SDS, 1 mM DTT, 10 μl/ml protease inhibitor) on ice for 30 min, followed by sonication (10 s, 20% amplification). The final solution contained the nuclear extract.

### Western blots

In brief, lysates were separated on polyacrylamide gels, transferred onto PVDF membranes, and blotted with the indicated antibodies: mouse monoclonal anti-Flag antibody (Abways, #AB0008), mouse monoclonal anti-ICP0 antibody (Santa, #sc-53070), mouse monoclonal anti-ICP8 antibody (Abcam, #ab20194), mouse monoclonal anti-VP16 antibody (Santa, #sc-7545), mouse monoclonal anti-β-actin antibody (Sino Biological, #1000166), anti-TK antibody (laboratory stock), mouse monoclonal anti-GAPDAH antibody (Abways, #AB0037), mouse monoclonal anti-Histone antibody (Sino Biological, #100005), rabbit polyclonal anti-dCas9 antibody (ABclonal, #A14997), goat anti-mouse IgG-HRP (Invitrogen, #31430), and goat anti-rabbit IgG (H+L)-HRP (Invitrogen, #32460).

### Immunofluorescence Staining

Cells were fixed in methyl alcohol at −80°C overnight, then permeated and blocked with PBS-TBH (0.1% Triton X-100 in 1XPBS, 10% FBS, and 1% bovine serum albumin (BSA) (Sigma, #WXBD5147V)), reacted with primary antibodies overnight at 4°C or 2 hr at RT, and secondary Alexa-Fluor-594-conjugated goat anti-rabbit (Invitrogen, #A11012) or Alexa-Fluor-488-conjugated goat anti-mouse (Invitrogen, #A32723) for 30 min at RT in dark. The slides were mounted with mounting medium and imaged with a Zeiss confocal microscope (ZEISS Imager Z20).

### sgRNA design

Genomic information of the HSV-1 (F) strain genome was downloaded from GenBank (Accession: GU734771.1). sgRNAs were designed using Cistrome (ht tp://cistrome.org/SSC/). Control sgRNA (5’-GGGGTAGGCGGAGCCTCAGG-3’) has no known targets in the human genome or HSV-1 genome.

### Electroporation

Cells were deattached with 0.1% trypsin, washed with DPBS, resuspended in 100 μl of electroporation buffer (Cell Line Nucleofector® Kit V, #VCA1003), mixed with DNA and/or RNA as indicated in each experiment and electroporated at 110 V for 10 ms by the electroporator (Lonza Amaxa Nucleofector 2B). After electroporation, the cells in each electroporation cup were mixed with 1 ml DMEM and placed back into a culture dish for subsequent experiments.

### *In vitro* transcription

I*n vitro* sgRNA transcription was performed using the Ribomax T7 large-scale RNA production kit (Promega, #P1300) according to the manufacturer’s instructions. In short, 2 to 4 μg of template DNA containing a T7 promoter was used for each reaction. The in vitro transcription mixture was incubated at 37°C for 3.5 hr, and RNase-free DNase I was added to remove the DNA template. The transcribed RNAs were treated with alkaline phosphatase (Thermo, #01137175) for 1 hr and cleaned up using an E.Z.N.A. miRNA Kit (Omega, #R6842-01). The final RNA stocks were quantified using a NanoDrop spectrophotometer, aliquoted and stored at −80°C.

### Confocal microscopy and image processing

Cell samples were imaged with a Nikon microscope (N-STORM) with 100 X-oil immersion lenses. The images were processed using NIS Elements software by time-lapse microscopy with Z-stacks. Imaris software (Bitplane) was used to perform 3D visualization and 3D reconstitution. DNA FISH spots were built using the Spots function, and the surface of the nucleus was built as a membrane object using the cell function and then switched to a surface object. The number of plasmids or HSV-BAC on the nuclear envelope was quantified by using the reconstituted plasmids or HSV-BAC spots and the nuclear periphery surface.

### Chromatin immunoprecipitation (ChIP)

ChIP assays were carried out as described previously. In short, at the indicated time points postinfection, cells were treated with 1% paraformaldehyde (Sigma, #P6148) for 10 min, washed twice with PBS, scraped off, resuspended in sodium dodecyl sulfate (SDS) lysis buffer (1% SDS 10 mM EDTA 50 mM Tris, pH 8.1) containing protease inhibitors (Thermo Scientific, #EO0492), and incubated on ice for 20 min. Cell lysates were sonicated in 20-s pulses for a total of 4 min to yield DNA fragments of 200 to 500 bp in length and further clarified by centrifugation at 13,000 g at 4°C for 10 min. The supernatant was collected and diluted 10-fold in phosphate-buffered radioimmunoprecipitation assay buffer (0.1% SDS, 1% sodium deoxycholate, 150 mM NaCl, 10 mM Na_2_PO_4_, 2 mM EDTA, 1% NP-40) and precleared for 1 hr at 4°C. At this point, 1% of the total volume was aliquoted as input. Immunoprecipitation was carried out at 4°C overnight with mouse immunoglobulin G (Cell Signaling Technology, 5415S) or anti-histone antibodies (anti-H3K4me3 antibody (Abcam, #ab8580), anti-H3K9me3 antibody (Abcam, #ab8898), anti-H3K27me3 antibody (Abcam, #ab6002), anti-H4K20me3 antibody (Abcam, #ab9053), anti-H3K4AC antibody (Abcam, #ab4441)). Immunocomplexes were collected by incubation with protein A/G beads (Santa Cruz, #sc-2003) for 1 hr at 4°C with rotation, washed sequentially with low-salt wash buffer (150 mM NaCl, 20 mM Tris HCl, pH 8.1, 2 mM EDTA, 1% Triton X-100, and 0.1% SDS), high-salt wash buffer (500 mM NaCl, 20 mM Tris HCl, pH 8.1, 2 mM EDTA, 1% Triton X-100, and 0.1% SDS), lithium chloride wash buffer (0.25 M LiCl, 1% NP-40, 1% sodium deoxycholate, 1 mM EDTA, and 10 mM Tris-HCl, pH 8.1), and Tris-EDTA buffer (10 mM Tris-HCl, pH 8, 1 mM EDTA) and eluted by incubation with elution buffer (1% SDS, 0.1 M NaHCO3) at RT for 10 min, at 65°C for 10 min, and finally at RT for 10 min. NaCl was added to both eluates and inputs to reach a final concentration of 0.2 M. All samples were then treated with RNase A (Omega, #D10SG) and proteinase K (QIAGEN, #160016374), and total DNA was purified by E.Z.N.A. Gel Extraction Kit (Omega, #D2500-02) and used as a template for qPCR.

The amounts of input and percentage of immunoprecipitated DNA were measured by quantitative PCR using primers specific for viral promoters and the cellular GAPDH pseudogene. The fraction of viral or host DNA immunoprecipitated with the relevant antibodies was normalized to their input values before further comparisons.

### Statistical analysis

In this study, each of the experiments was performed with at least three biological replicates unless otherwise specified. Data are presented as the mean ± sd. calculated by GraphPad Prism 6.0 software. Two-tailed unpaired Student’s t-test, ordinary one-way or two-way ANOVA indicated in each figure were used to calculate P values. N. S represents not significant, P > 0.05, “*” represents P ≤ 0.05, “**” represents P ≤ 0.01, “***” represents P ≤ 0.001, and “****” represents P ≤ 0.0001.

## Results

### A two-component CRISPR-nuPin platform that instant nuclear edging of extrachromosomal DNAs

The life cycle of many DNA viruses measures in the unit of hours [32], and thus, the probing tool to interrogate the spatial interactions between viral genomes and the host subnuclear environment needs to act quickly. To simplify the DNA repositioning system [33–35] and avoid the possible time lag introduced by a second layer of regulation (ABI-PYL1 or CIBN-CRY2), we hypothesized that a two-component strategy, a subnuclear compartment tethered to dcas9 and a sgRNA introduced by electroporation, was able to mediate inducible and fleet relocation of viral DNA genomes (Figure 1 A). To achieve nuclear periphery docking, Flag-emerin-TEV (TEV protease recognition sequence)-GFP was fused to the C-terminus of *Streptococcus pyogenes* dCas9 (D10A&H840A) (dCas9-emerin), resulting in a fusion protein with a predicted size of 230 kDa (Sup Figure 1 A). Insertion of a nuclear localization signal (NLS) at the N- and C-termini of dCas9 enabled nuclear importation of the majority of dCas9-emerin, while Flag-tagged emerin localized both in the nucleus and cytoplasm (Figure 1 B and Supplementary Figure 1 A). We then constructed a doxycycline (DOX)-inducible dCas9-emerin-expressing HEp-2 cell line (dCas9-emerin) using a lentiviral transduction system. A HEp-2 cell line expressing only dcas9 protein was established simultaneously and served as a control cell line (dCas9). Successful induction of the dCas9-emerin fusion protein in dCas9-emerin cells by DOX treatment was confirmed by immunoblotting. The colocalization of anti-Flag and anti-dCas9 antibodies recognized epitopes along the outer edge of the DAPI-stained nuclear area under an immunofluorescence microscope, indicating that the dcas9-emerin fusion protein properly localized to the inner nuclear membrane (Figure 1 C, Sup Figure 1 B, C). DOX treatment had no detectable cytotoxicity on HEp-2 cells (Sup Figure 1 D).

To test the time efficiency of the system, dCas9-emerin cells were first electroporated with a 5 kb plasmid (Sup Figure 1 G, H) or a 170 kb BAC DNA (Figure 1 D, E) and again with its targeting sgRNAs at 24 hr post initial electroporation. Representative 3D images of the nucleus at the indicated time points post sgRNA electroporation (upper panel) and their 3D reconstruction images (bottom panel) are shown (Figure 1 D). Targeted DNAs were instantly repositioned to the nuclear margin upon induction of sgRNAs, and more than 60% of BAC DNA was relocated away from the inner nuclear space as soon as 10 seconds post sgRNA electroporation (Figure 1 E). To test the repositioning efficiency of CRISPR-nuPin on the host genomic locus, dCas9-emerin cells were electroporated with sgRNA targeting the telomere region (Figure 1 F, G). At 3 hr post sgRNA induction, more than 53% of cellular telomeres were detected bound to the nuclear edge compared to approximately 27% at time 0 hr. Our results indicate that the CRISPR-nuPin system is capable of instant repositioning of extrachromosomal DNA of up to 170 kb in size to the nuclear periphery. Note that a single dot in the 3D reconstruction images represents single or multiple plasmids or BACs.

### HSV-1 genomes positioned to the nuclear margin were subjected to transcriptional repression

To initiate replication, DNA viruses need to eject their genetic material into the host nuclei and initiate viral gene transcription. It is conceivable that individual viral genomes encounter highly heterogenic microenvironments upon entry. Indeed, spatially, approximately 25% of the HSV-1 genomes localized to the edge of the inner nuclear membrane during natural infection, and the remaining viral DNA penetrated further into the nuclear interior but was still within proximity to the nuclear periphery (Sup Figure 2 A). We asked if viral DNAs had a preference for the inner nuclear space over the nuclear edge for replication. To this end, 5 sgRNAs of different numbers of targeting sites within the HSV-1 genome were designed to mediate nuclear editing of HSV-1 genomes in the CRISPR-nuPin system. Cautions were made to avoid protein coding regions of the virus (Sup Figure 2 B). All sgRNAs were able to mediate significant inhibition of HSV-1 in dCas9-emerin cells (Sup Figure 2 C). Since sgRNA2 binds to one site at the region coding for the 8.3 kb latency-associated transcript (LAT) precursor of HSV-1, which plays no essential role during HSV-1 lytic replication, and another site in the terminal repeat short regions (TRs), it was used to mediate HSV-1 genome repositioning in the following experiments in this study. To exclude the effect of nonspecific factors such as dCas9-sgRNA binding to viral genomes or sgRNA-induced innate immunity responses, HEp-2 cells expressing GFP, dCas9 or dCas9-emerin fusion protein were transfected with a plasmid encoding a non-HSV-1 targeting sgRNA (Ctr sgRNA) or the HSV-1 targeting sgRNA2 (HSV-1 sgRNA). These cells showed no essential differences in supporting HSV-1 replication, except for those expressing both the dCas9-emerin fusion protein and HSV-1 sgRNA (Sup Figure 2 D, E, F). The single-cycle growth kinetics of HSV-1 were measured in dCas9-emerin cells infected at an MOI of 1 in the presence of Ctr sgRNA or HSV-1 sgRNA. In dCas9-emerin cells expressing HSV-1 sgRNA, viral genomes were guided to the nuclear edge compared to the Ctr sgRNA group (Figure 2 A a, b), and tethering viral genomes to the inner nuclear membrane led to reduced virus production at all time points, including the plateau phase (Figure 2 A). Increasing the multiplicity of infection up to 20 infectious virions per cell did not help the virus overcome the plight (Figure 2 B, Sup Figure 2 G).

HSV-1 transcribes three sets of viral genes in chronological order [36–38]. Intriguingly, in the host nuclei with the viral genomes positioned to the nuclear edge, the mRNA levels of the representative viral genes of the three classes declined sharply to a significantly lower basal level before transcription of these viral genes was fully activated. Although the launch of full-speed transcription of ICP27, a representative α gene of HSV-1, in HSV-1 sgRNA-transfected dCas9-emerin cells was delayed by 30 minutes compared to that in the control group, the transcription rate, once switched to the higher gear, was comparable. The total amount of ICP27 mRNA synthesized by nuclear-edged HSV-1 remained 10-fold lower than that synthesized by HSV-1 freely infecting the nuclei (Figure 2 C). Similar patterns were observed for representative β (TK) and γ (VP16) genes with a slightly longer delay of full-speed transcriptional onset (Figure 2 C). Postponed transcriptional onsets led to delayed accumulation of viral proteins such as ICP8, ICP0, VP16 and TK (Figure 2 D) and ineffective replication center formation efficiency (Sup Figure 3 A) [31]. The observations in this set of experiments showed that deposition of HSV-1 genomes to the nuclear border area led to decreased initial basal transcription and postponed full-speed transcription onset of the viral gene cascade, severely delayed viral protein accumulation and ultimately inefficient replication of the virus.

The nuclear periphery is a known area decorated with heterochromatin. To investigate whether viral genomes at the nuclear edge were subjected to histone-mediated transcriptional suppression, ChIP-qPCR quantifying the association level of actively and repressively modified histones to the promoter regions of HSV-1 α, β and γ genes was conducted. For viral genomes positioned to the nuclear edge, promoters of the representative viral genes showed significantly less association with active histones (H3K4me3 and H3K9ac) and increased frequency with repressive histones (H4K20me3, H3K9me3, and H3K27me3) compared with those unmanipulated viral DNAs in the nuclei (Figure 2 E). Furthermore, the repressive histone packaging, represented by H3K9me3, of viral genomes at the nuclear edge progressively increased over time during the first hour of infection (Figure 2 F). Whole-cell inhibition of histone deacetylase (HDAC) activity by treatment with sodium butyrate (SBt) relieved the transcriptional suppression of the nuclear-edged viral genomes to some extent but did not completely complement the nuclear margining-imposed transcriptional disadvantage on HSV-1 (Figure 2 G).

These results, in all, lead to an intriguing inference that intranuclear viral genomes face diversified fates upon nuclear entry and that those wandering in the nuclear edge area are underprivileged during host-virus competition during lytic infection.

### Relocation of replicating HSV-1 genomes at 1 hpi significantly promoted transcription of viral β and γ genes, termed an “Escaping” effect

HSV-1 genomes entering the nuclei naked and will encounter waves of restrictive responses imposed by the host, some of which may also be taken advantage of by the virus [39, 40]. In this section, experiments were conducted in an attempt to establish the chronology of host responses to incoming viral genomes. To this end, dCas9-emerin cells infected with HSV-1 at an MOI of 1 were electroporated with HSV-1 sgRNA or Ctr sgRNA at 0 (immediately after 2 hr of virus inoculation on ice), 0.5, 1 and 2 hpi (Figure 3 A). Examination of the subnuclear localizations of HSV-1 genomes at 5 minutes post sgRNA electroporation by FISH showed that the CRISPR-nuPin system mediated prompt and highly effective editing of viral genomes in the nucleus during active virus infection (Figure 3 B). Titration of the virus yields in cells with HSV-1 genomes disturbed and repositioned to the nuclear border at different time points during early infection showed that the inhibitory effect imposed by nuclear edging of HSV-1 genomes was time sensitive and was only effective during the very early and short period of time (approximately 0.5 to 1 hr under this experimental condition) (Figure 3 C). An appealing phenomenon was observed when HSV-1 genomes were dislodged from their initial intranuclear position at 1 hpi, and the virus yield was significantly boosted (Figure 3 C). Tracking virus production at multiple time points during infection confirmed that relocation of HSV-1 genomes to the nuclear periphery at 1 hpi switched the virus production efficiency to a significantly higher gear as early as 4 hpi (Figure 3 D). We thus refer to the promotive replication mediated by relocation of HSV-1 genomes away from their original spots at 1 to 1.5 hpi as an “Escaping” event.

**Figure 3.**
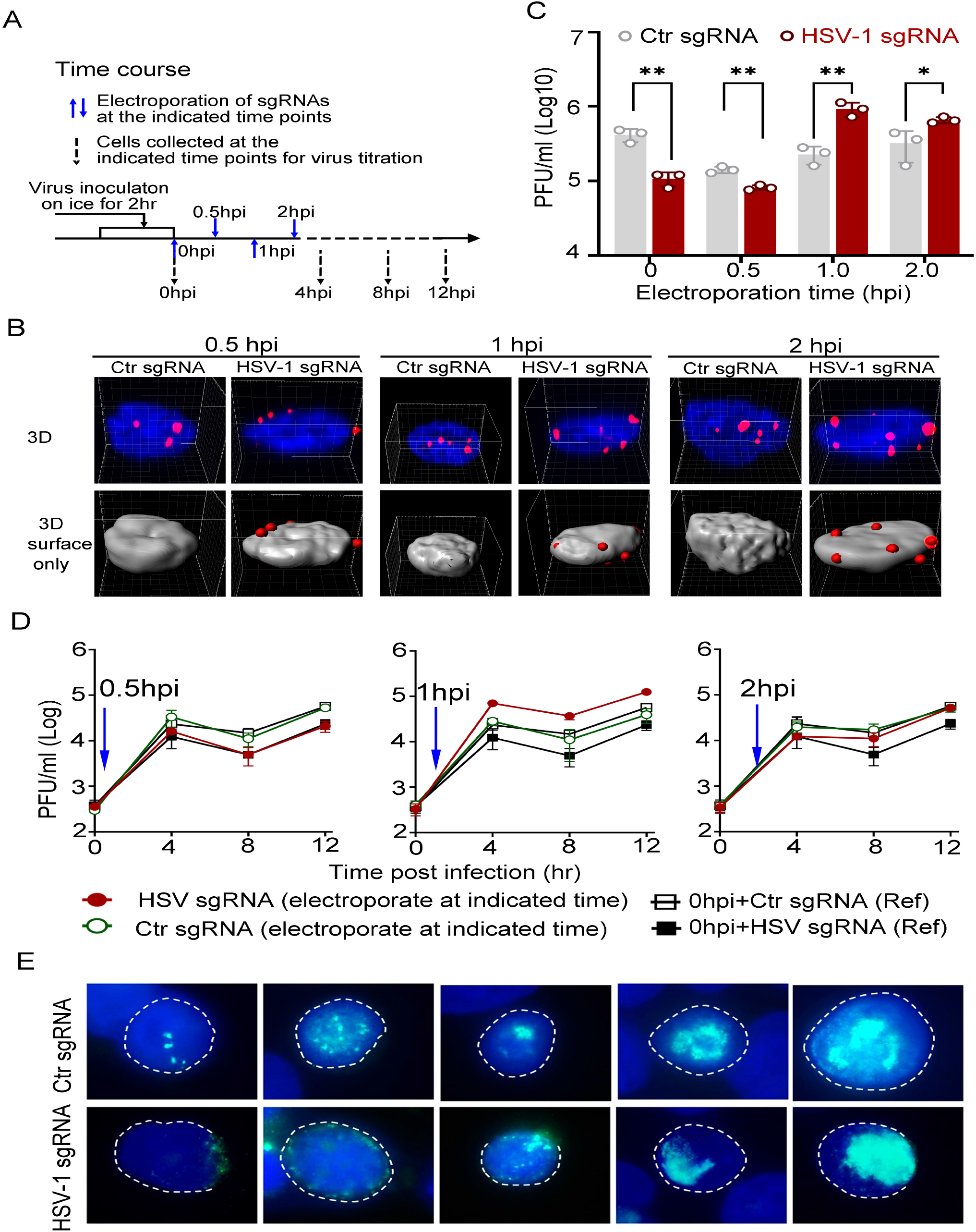
Dislodging HSV-1 genomes to the nuclear edge at 1 hpi promoted virus infection, termed “Escaping”. **A.** Schematics of the experiments. dCas9-emerin cells were infected with HSV-1 virus at an MOI of 1 and electroporated with HSV-1 sgRNA or Ctr sgRNA at 0 (immediately post virus inoculation), 0.5, 1 and 2 hpi. **B.** At 5 min after electroporation, cells were fixed, and intranuclear HSV-1 genomes were stained by FISH (red) and scanned under a Nikon microscope (N-STORM). Representative Z-series images are shown in the upper panel, and 3D reconstruction of the images showing only surface staining is shown in the bottom panel. **C.** At 12 hpi, cell-associated virus was titrated by plaque assay. **D.** Cell-associated virus yields were tracked for 12 hr post infection by plaque assay. dCas9-emerin cells electroporated with Ctr sgRNA (empty rectangle) or HSV-1 sgRNA (filled black rectangle) at 0 hpi served as a reference and were plotted repeatedly in three charts. The blue arrow indicates the time of sgRNA electroporation. The red filled cycle represents the virus yield in cells that received HSV-1 sgRNA, and the empty green cycle represents the virus yield in cells that received Ctr sgRNA. **E.** At 4 hpi, cells were fixed and stained with anti-ICP8 (green). Representative images of all stages of the replication center of HSV-1 are shown.

We asked if nuclear editing of HSV-1 genomes at 1 hpi facilitated replication assembly of the virus. HSV-1-infected cells that received viral-specific sgRNA or Ctr sgRNA at 1 hpi were examined for viral replication center formation efficiency at 6 hpi [31]. As shown in Figure 3 E, all stages of replication centers, delegated by aggregation of the viral single-stranded DNA binding protein ICP8, were detected in both HSV-1 genome-edged and naturally dispersed groups with no distinguishable overall pattern differences, indicating that the assembly of viral replication centers occurs normally at the nuclear margin (Figure 3 E).

### Early accumulation of ICP0 is a sufficient but not necessary condition of “Escaping”

To obtain a more integrative picture, the expression patterns of viral genes and the accumulation of viral proteins were examined for viruses in the “Escaping” event, and we reported the following observations: (For the sake of simplicity, “Electroporation of HSV-1 sgRNA at 1 hpi during HSV-1 infection of dCas9-emerin cells” was abbreviated as “E1hpi”). I) E1hpi led to elevated expression of ICP27 and ICP0 (α genes) starting from 4 to 5 hpi and significantly boosted transcription of TK (β gene) and VP16 (γ gene) starting from 3 hpi (Figure 4 A). II) E1hpi led to elevated viral protein levels of ICP0, ICP8, TK, and VP16 at 8hpi and 12hpi (Figure 4 B). III) Early accumulation of ICP0 protein between 3 and 4 hpi in the E1hpi group was detected in the absence of mRNA level change (Figure 4 C). In summary, we found that dislodging viral genomes from their previously occupied niches during a specific time window amid infection (1-1.5 hpi) led to stabilization of ICP0 and boosted viral gene expression.

**Figure 4.**
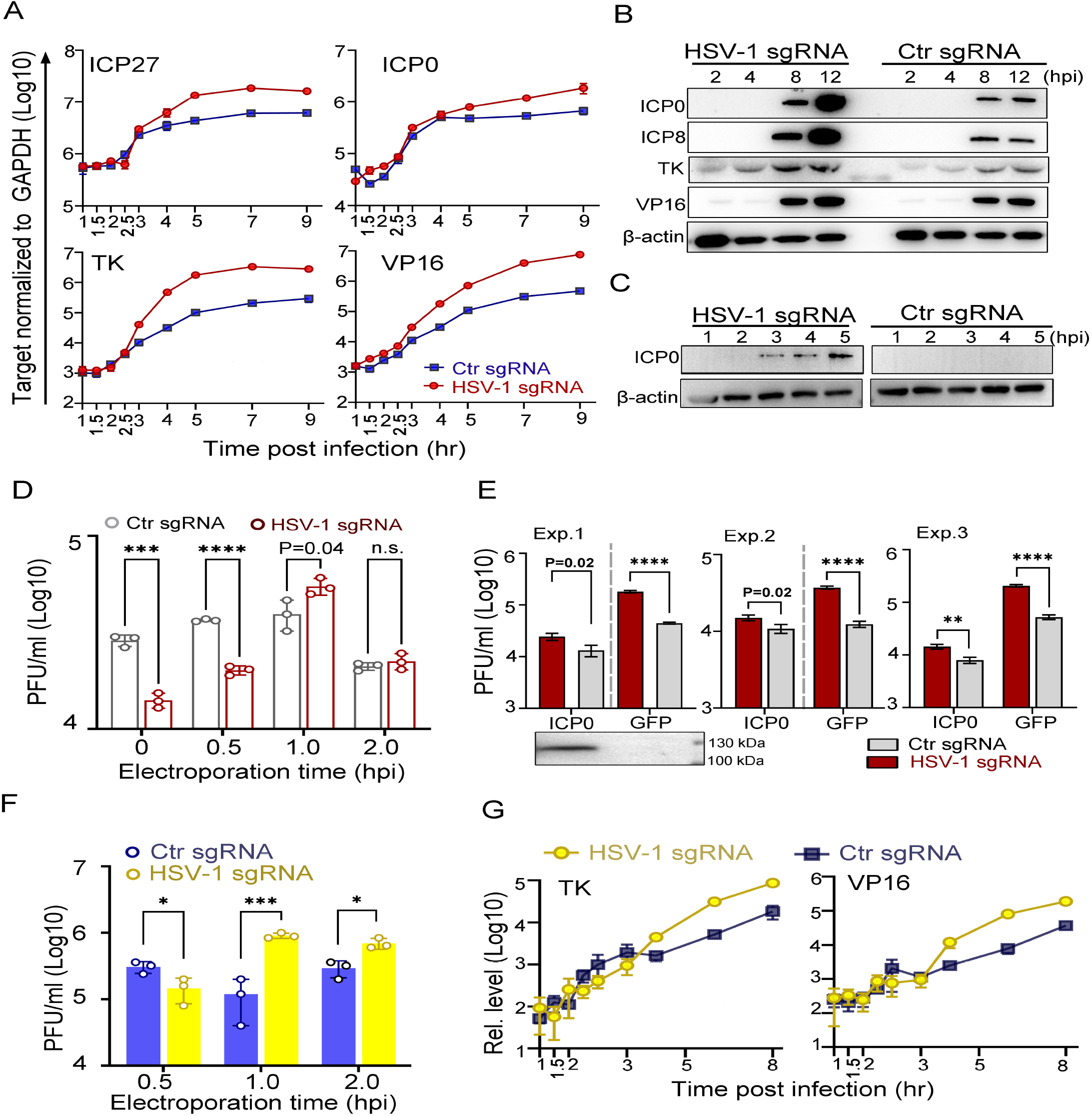
ICP0 is a sufficient but not necessary condition of “Escaping”. **A-C.** dCas9-emerin cells infected with HSV-1 virus at an MOI of 1 were electroporated with HSV-1 sgRNA or Ctr sgRNA at 1 hpi. **A.** The mRNA levels of ICP27, TK, ICP0 and VP16 at the indicated time points were measured by qPCR. **B.C.** Protein levels of ICP0, ICP8, TK and VP16 were measured by immunoblot. Five-fold more total protein was loaded in the C ICP0 panel to detect trace amounts of the protein. **D-E.** dCas9-emerin cells expressing ICP0 or GFP were infected with HSV-1 at an MOI of 5 and electroporated with HSV-1 sgRNA or Ctr sgRNA at 0, 0.5, 1 and 2 hpi (D) or at 1 hpi (E). The HSV-1 virus yield at 12 hpi was titrated and plotted. Expression of ICP0 was confirmed by immunoblotting. F. A similar experiment as in D was performed with ΔICP0 virus. G. A similar experiment as in A was performed with ΔICP0 virus.

As ICP0 is an essential transactivator that regulates the transcription of the β and γ genes, we investigated the necessity and sufficiency of ICP0 in this “escaping” event. Prefilling of host nuclei with ICP0 protein mitigated the intranuclear spatial bias of HSV-1, although placing incoming viral genomes to the nuclear edge still posed a significant disadvantage on its growth (Figure 3 D and Figure 4 D). Importantly, “Escaping” in ICP0-overexpressing cells mediated only a modest to nonpromotive effect on virus replication (P=0.04) (Figure 4 D). To further confirm the role of ICP0, the “escaping” effect in dCas9-emerin cells transfected with plasmids coding for GFP or HSV-1 ICP0 was compared. While “Escaping” promoted virus growth by approximately 4-fold in GFP-overexpressing cells, it boosted HSV-1 production by only 2-fold in ICP0-overexpressing cells in three independently repeated experiments (Figure 4 E), implying that early accumulation of ICP0 protein was a contributing factor of the “Escaping” effect. To test whether ICP0 was the sole factor, a mutant HSV-1 virus lacking ICP0 (ΔICP0) was used for infection. ΔICP0 virus was similarly susceptible to nuclear editing-mediated suppression of incoming viral genomes (Sup Figure 4), and “Escaping” significantly facilitated viral growth and promoted transcription of viral β and γ genes of ΔICP0 (Figure 4 F, G), suggesting that ICP0 is not a necessity during “Escaping”.

In summary, the observed “Escaping” phenomenon hinted that a wave of inhibitory factors was loaded onto HSV-1 genomes at 1 to 1.5 hpi to restrict the expression of viral genes, concurrent with ICP0 destabilization.

## Discussion

### CRISPR-nuPin is an inducible two-component platform that mediates rapid repositioning of extrachromosomal DNAs to the nuclear edge

Previously reported CRISPR-based genome re-organization systems use chemical- or light-inducible approaches to mediate large-scale spatial genome reorganization/engineering, and the average reported response time for these systems to relocate genomic DNA to the nuclear periphery is in the range of hours [33–35, 41, 42]. To increase simplicity, reduce the possible time lag resulting from chemical addition and/or diffusion and manipulate the subnuclear localization of viral genomes amid infection instantly, we developed a two-component CRISPR-nuPin strategy that mediated rapid relocation of extrachromosomal DNAs up to 170 kb within minutes. CRISPR-nuPin utilized a dCas9-emerin fusion protein to tether targeted DNAs to the inner nuclear envelope upon electroporation of sgRNAs. The CRISPR-nuPin strategy is advantageous in the following aspects: I) CRIPSR-nuPin is induced through electroporation of sgRNAs and mediates prompt repositioning of targeted DNAs to the nuclear edge and is applicable to varying subcellular docking sites. II) CRISPR-nuPin operates in live cells and works efficiently and fleetly during active virus infection, making it a valuable tool to interrogate the complex virus and host interactions in space and time. III) CRISPR-nuPin is in principle applicable to various important DNA viruses, providing an unprecedented tactic to probe subcellular viral infection at the spatial-temporal scales. A typical example reported herein used CRISPR-nuPin to synchronize and investigate the 3D positions of viral genomes and their virological fates.

### Nuclear periphery: an edge of darkness for DNA viruses?

The nuclear periphery is conventionally considered a region of inactive chromatin, and nuclear envelope association frequently correlates with transcriptional repression of endogenous genes. However, artificially tethering genomic loci to the inner nuclear membrane has been shown to suppress transcription from the repositioned regions only moderately or not at all. This genomic locus-specific sensitivity to nuclear periphery repositioning implies that complex mechanisms in addition to physical intranuclear localizations regulate the expression of endogenous genes [43–51]. The study in this report showed that approximately 25-30% of HSV-1 genomes localize at the nuclear edge during natural infection in HEp-2 cells (Sup Figure 2 A). It is then a tantalizing question whether the nuclear edge represses transcription from the incoming viral DNAs. As shown in this report, HSV-1 genomes experienced a short period of basal transcription before the full on-set of α gene transcription, which in the case of HSV-1 infecting HEp-2 cells at an MOI of 5 was approximately 30 minutes post virus initial attachment to the cell surface (Figure 2 C). Tethering viral genomes to the inner nuclear envelope during this time-window resulted in increased association with repressive histones, stronger transcriptional suppression, and prominently reduced virus yields. This provided the first experimental evidence showing that the nuclear edge imposed more stringent transcription silencing on the incoming viral DNAs and that HSV-1 viral genomes reaching further into the nuclear interior upon entry were poised for more efficient transcription and replication.

We referred to this initial “nuclear edging” susceptible phase as the nuclear periphery sensitive phase (Figure 5). Of note, prior studies have shown that HSV-1 relies on A-type lamin to avoid heterochromatin packaging and to efficiently organize VP16 activator complex formation to activate viral α genes and that small HSV-1 replication centers are frequently found localized along the nuclear periphery [52–54]. The apparent contradictions could be attributed to I) the intrinsic systemic differences between the investigation platforms on which these studies were performed. Previous studies were conducted in A-type lamin-deficient mouse embryo fibroblasts, which show massive abnormities in nuclear envelope integrity, genome organization, heterochromatin distribution, etc. [55, 56]. This study used HEp-2 cells, a cell line commonly used for HSV-1 virus propagation and research. II) The difference in the scope of physical distance referred to by “the nuclear edge” and “the nuclear periphery” in these studies. III) The timeframe post infection that these studies are investigating.

**Figure 5.**
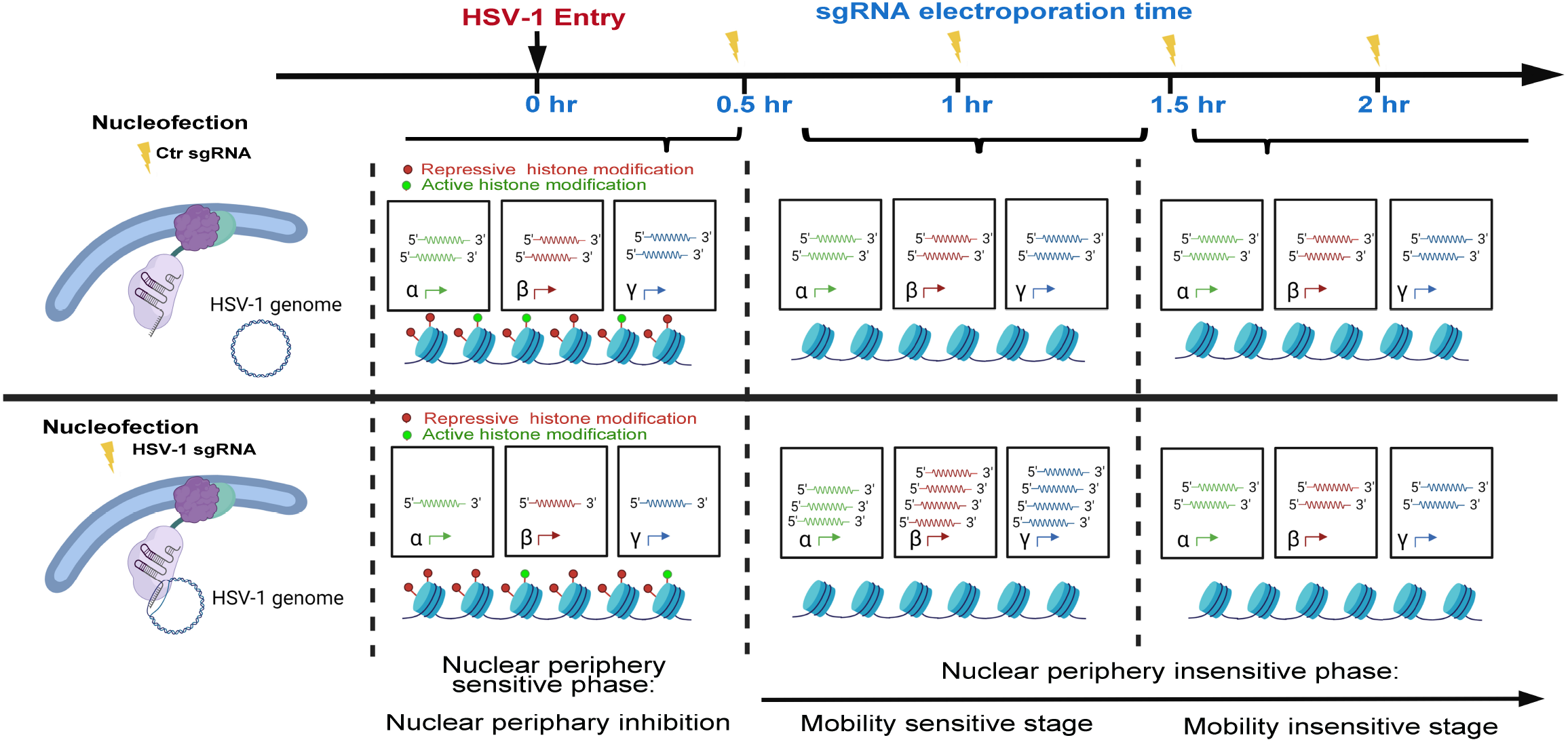
Proposed stages of the early intranuclear life of HSV-1.

### Spatial-temporal interactions between HSV-1 and host factors: fierce competition with heterogeneity

In HEp-2 cells infected with HSV-1 at an MOI of 1, HSV-1 genomes were poised for efficient infection and no longer susceptible to nuclear peripheral positioning-mediated suppression after 1 hpi. We referred to this period as the second phase of early infection—the nuclear periphery-insensitive phase (Figure 5). Interestingly, there was a short time window when dragging HSV-1 genomes away from their previously occupied spots by CRISPR-nuPin significantly promoted transcription from viral DNAs, especially for the β and γ genes, suggesting that an increased mobility/turbulence at approximately 1 to 2 hpi may help the viral genomes escape the ensuing suppressions. We thus refer to 0.5 to 1.5 hpi as the inhibitive stage and infer that during the inhibitive stage, the viral genomes with increased movability may gain growth advantages. After the inhibitive stage, intranuclear repositioning of viral genomes no longer caused detectable changes in the final virus yield (2 hpi in HEp-2 cells infected with HSV-1 at an MOI of 1), indicating that the virus pulled off the first virus-host competition at this time point (Figure 5).

In summary, this example study using CRISPR-nuPin on HSV-1 early life in the nucleus has shown that not only do viral genomes entering the nucleus confront heterogeneous microenvironments, but increased mobility or turbulence of viral genomes during infection may contribute to viral pathogenic outcomes.

## Acknowledgement

This project is supported by the National Natural Science Foundation of China (No. 31870157 and 81802006) and the Shenzhen Science and Technology Innovation Program (JCYJ20180307151536743 and KQTD20180411143323605).

## Declaration of interest statement

The authors declare no conflicts of interest.

## Supplementary Figures

**Figure 1.**
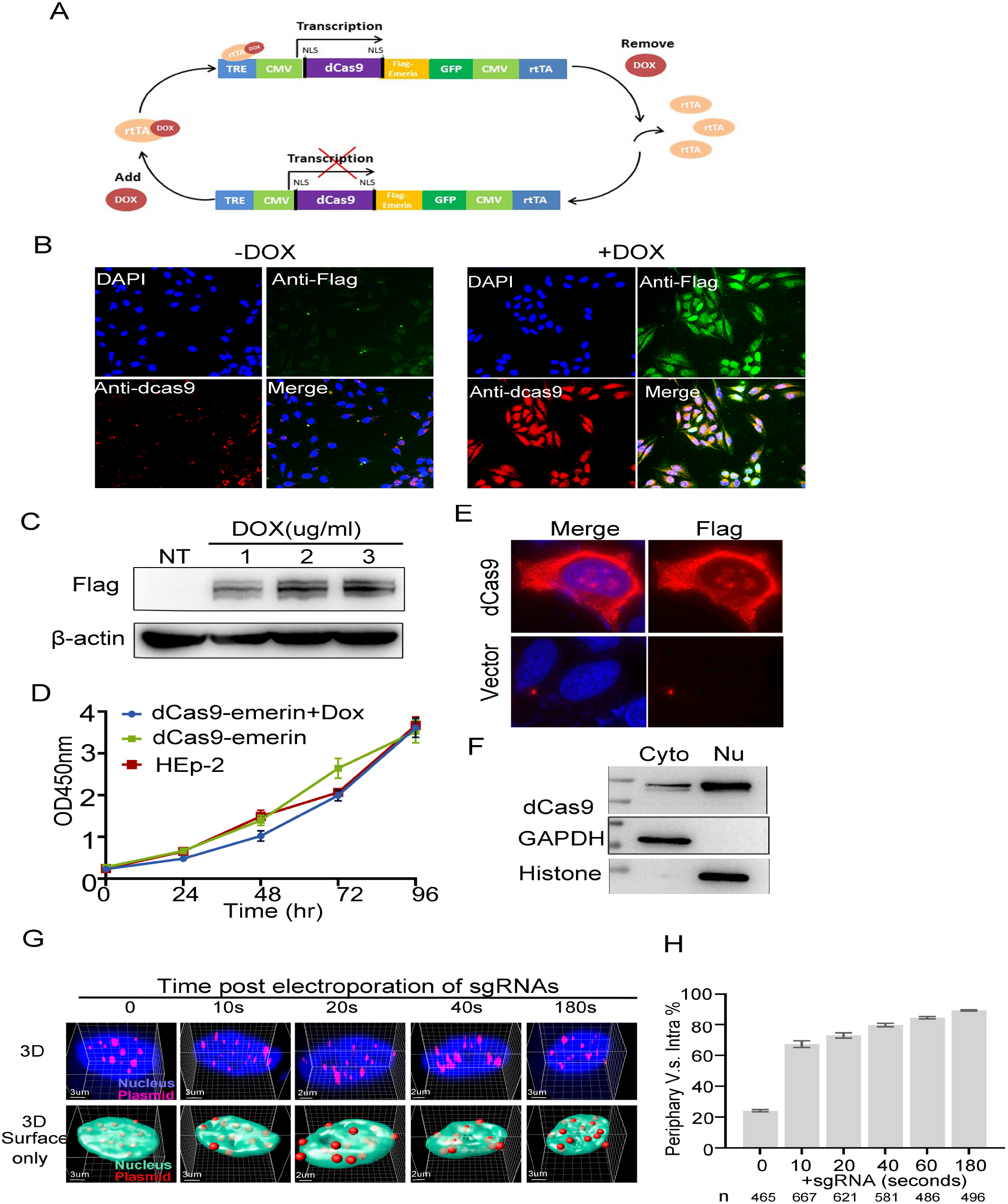
General characterization of the CRISPR-nuPin system. **A.** Schematic of the DOX-induced CRISPR-nuPin expression cassette in dCas9-emerin cells. **B.** Percentage of dCas9-emerin-expressing cells in the dCas9-emerin cell line. dCas9-emerin cells were treated with DOX (+DOX) or DMSO (-DOX) and stained with antibodies against Flag (green) and dcas9 (red). **C.** dCas9-emerin cells were treated with DOX at the indicated concentration for 48 hr, and the expression of dCas9-emerin was examined by immunoblotting. **D.** HEp-2 and dCas9-emerin cells treated with DOX (+DOX) or DMSO (-DOX) (1000) were seeded onto 96-well plates. Cell viability was tested using CCK-8 reagent for 96 hr. **E, F.** Confirmation of dCas9 protein expression in transiently transfected HEp-2 cells by immunofluorescence staining (E) and immunoblotting of subcellular fractions with anti-dCas9 antibody (F). **G, H.** dCas9-emerin cells (+DOX) transfected with 5 kb plasmids were electroporated with the plasmid-specific sgRNA. At the indicated time points post electroporation, cells were fixed, and the 5 kb plasmids were stained by FISH and scanned under a Nikon microscope (N-STORM). Representative Z series images are shown in the upper panel of G, and 3D reconstruction of the images is displayed in the bottom panel. The percentage of the plasmid stains located at the nuclear edge versus total intranuclear stains at each time point was calculated and plotted. Total counted stains at each time point are shown as n in the bottom of H.

**Figure 2.**
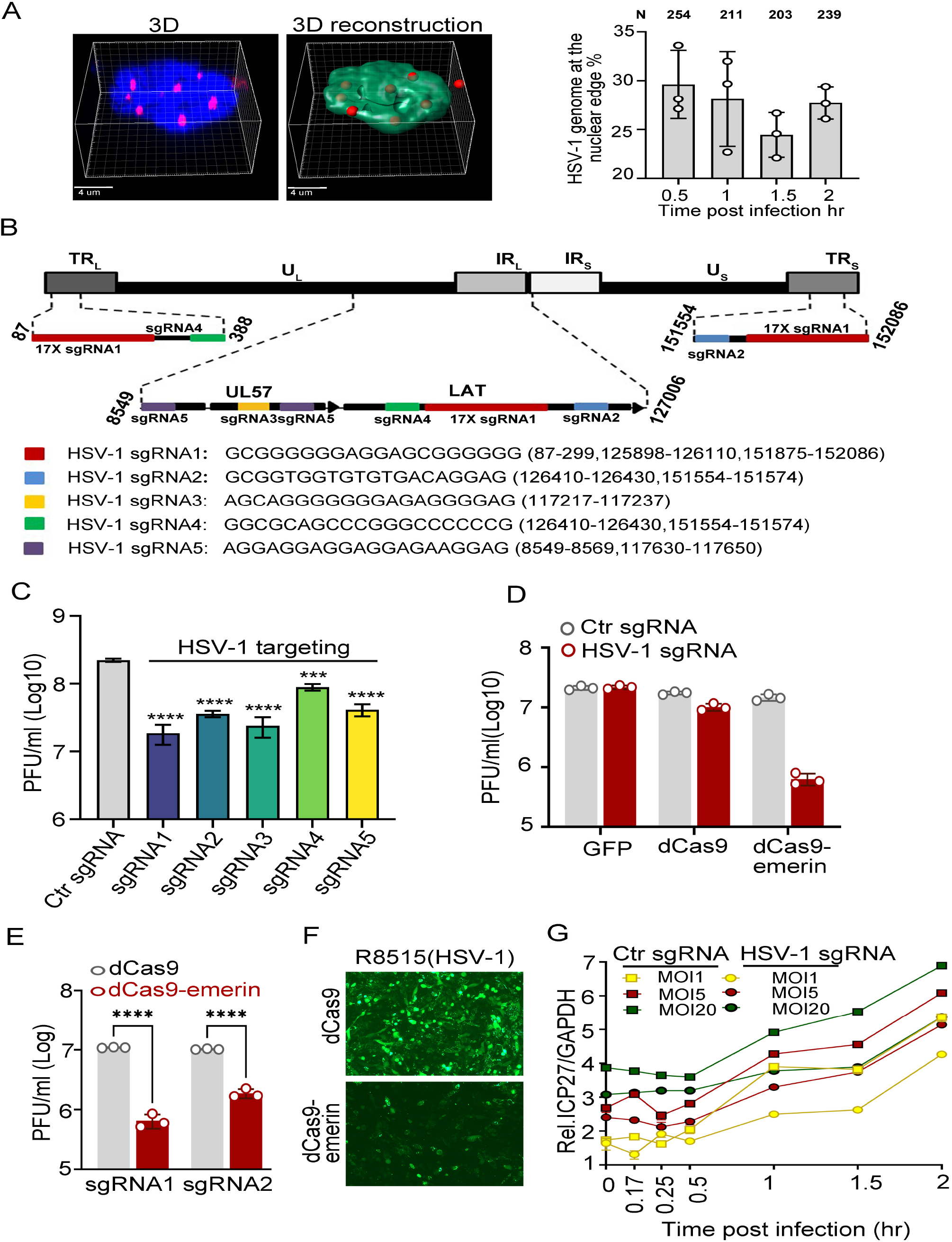
HSV-1 genomes were diversely dispersed in the nucleus upon entry, and editing of viral genomes suppressed virus replication. **A.** HEp-2 cells were infected with HSV-1 at an MOI of 5. Cells were fixed at 0.5, 1, 1.5 and 2 hpi, and the intranuclear HSV-1 genomes were stained by FISH and scanned under a Nikon microscope (N-STORM). Representative Z series image at 2 hpi and its 3D reconstruction showing only the surface of the DAPI-stained area are shown on the left. HSV-1 genomes localized to the nuclear edge versus total intranuclear viral genomes were counted and plotted on the right. N represents the number of viral genomes counted. **B.** Positions and number of targets of 5 HSV-1 sgRNAs in the viral genome. **C.** dCas9-emerin cells expressing Ctr sgRNA or each of the 5 HSV-1 sgRNAs were infected with HSV-1 at an MOI of 5. Virus yield at 24 hpi was quantified by plaque assay. **D.** A stable GFP-expressing HEp-2 cell line (GFP) and dCas9 and dCas9-emerin cells were transfected with plasmids expressing HSV-1 sgRNA2 or Ctr sgRNA for 24 hr and infected with HSV-1 at an MOI of 1. Cell-associated virus at 24 hpi was titrated by plaque assay. **E.** dCas9 and dCas9-emerin cells were transfected with plasmids expressing HSV-1 sgRNA1, sgRNA2 or Ctr sgRNA for 24 hr and infected with HSV-1 at an MOI of 1. Cell-associated virus at 24 hpi was titrated by plaque assay. **F.** dCas9 and dCas9-emerin cells were transfected with plasmids expressing HSV-1 sgRNA2 for 24 hr and infected with a recombinant GFP-expressing HSV-1 virus (R8515) at an MOI of 1. Cells were photographed at 24 hpi, and representative images are shown. **G.** dCas9-emerin cells transfected with plasmids expressing Ctr sgRNA or HSV-1 sgRNA2 for 24 hr were infected with HSV-1 at different MOIs. The mRNA level of ICP27 at the indicated time points post infection was measured by qPCR.

**Figure 3.**
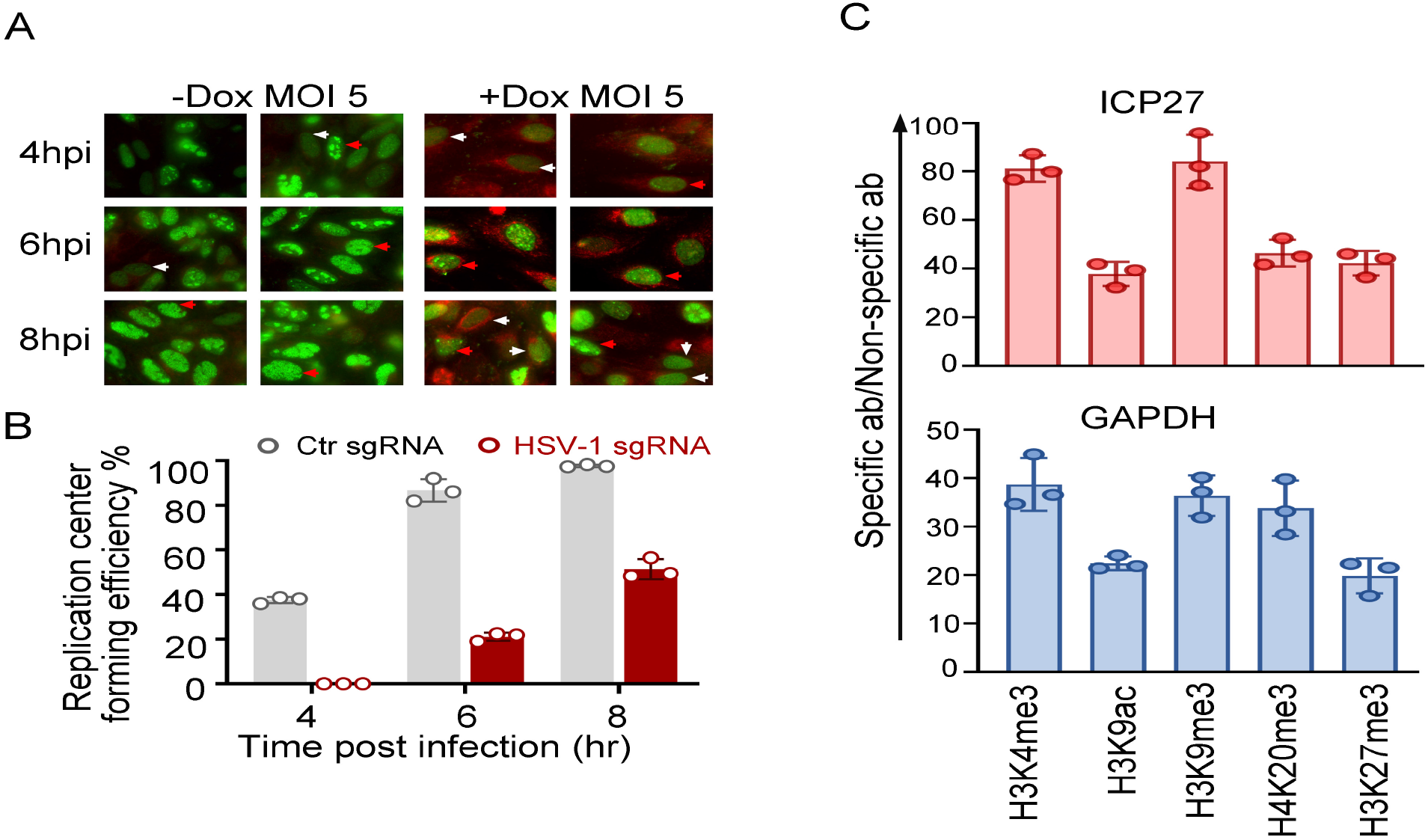
Replication of HSV-1 was affected in dCas9-emerin cells with the viral genomes positioned to the nuclear edge upon entry and quality controls of ChIP analyses. **A-B.** dCas9-emerin cells transfected with plasmids expressing HSV-1 sgRNA or Ctr sgRNA for 24 hr were infected with HSV-1 at an MOI of 5. At the indicated time point post infection, cells were fixed and stained with anti-ICP8 (green) and anti-Flag (red) antibodies. **A.** Two representative images of each time point are shown. White arrows indicate nuclei with diffuse ICP8 (inefficient replication center formation), and red arrows indicate nuclei with aggregated ICP8 (replication center formation). **B.** More than 200 nuclei in each sample were counted, and the HSV-1 replication center formation efficiency (nuclei of aggregated ICP8 versus total ICP8-positive nuclei) was quantified and plotted. **C.** The fraction of viral DNA (ICP27) (top panel) or host genomic DNA (GAPDH) (bottom panel) immunoprecipitated with the indicated antibodies was compared to their respective fraction immunoprecipitated by a nonspecific antibody (ab) (IgG) and plotted.

**Figure 4.**
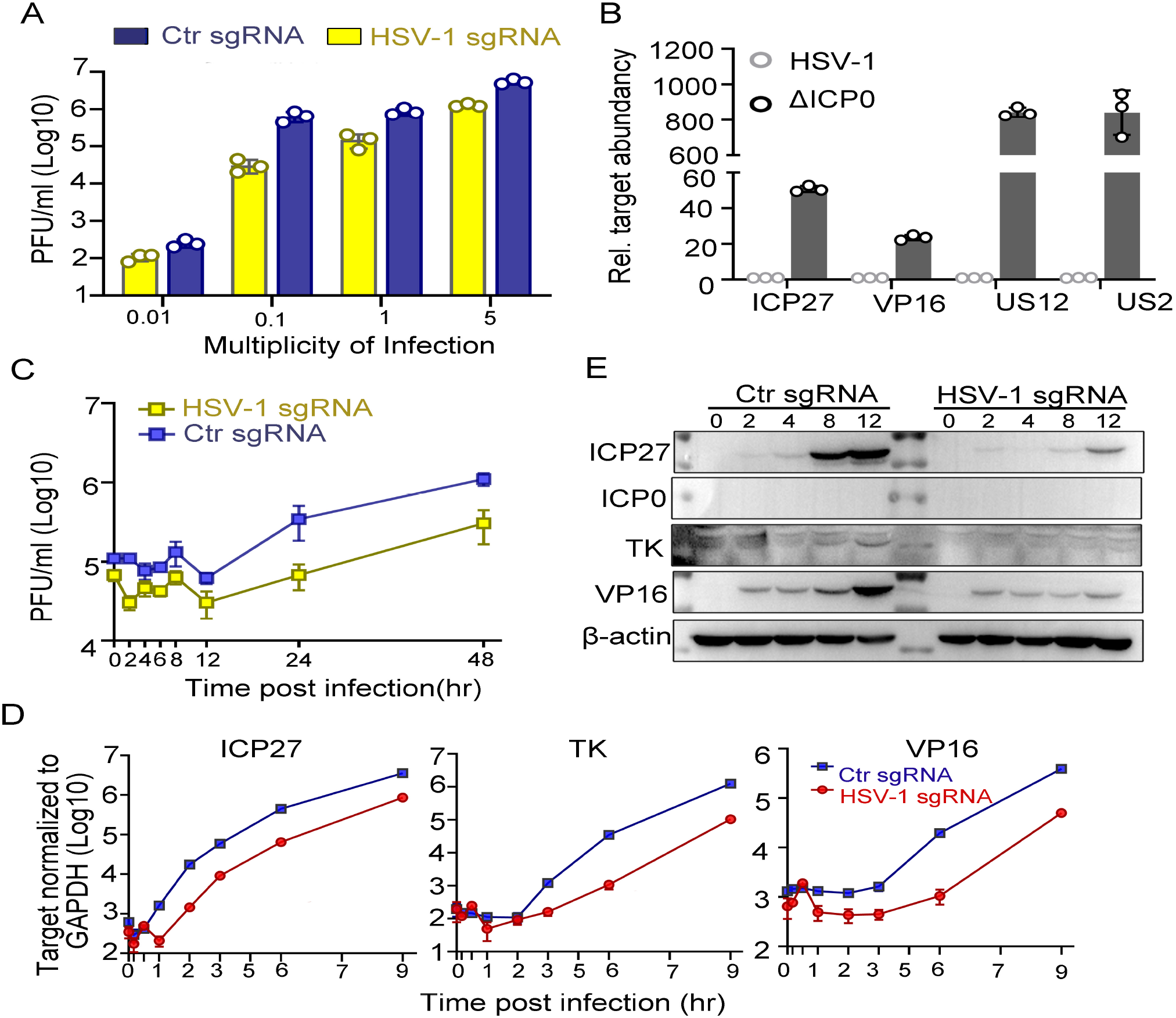
Characterization of the growth of ΔICP0 in dCas9-emerin cells with viral genomes inserted into the nucleus upon their entry. dCas9-emerin cells expressing HSV-1 sgRNA or Ctr sgRNA for 24 hr were infected with ΔICP0 virus at different MOIs (A) or an MOI of 1 (C-E). **A. The** ΔICP0 virus titer was determined by plaque assay at 12 hpi. B. Viral DNA was extracted from virus stocks of ΔICP0 or HSV-1 containing the same number of infectious virions (6×10^6^). The relative copy number of the indicated viral genes was measured by qPCR, and the readings of the ΔICP0 virus stock were compared to those of the HSV-1 virus stock (set as 1). **C.** ΔICP0 virus yield at different times post infection was measured by plaque assay. **D.** Accumulation of viral mRNA (ICP27, TK and VP16) during infection was quantified by qPCR. **E.** Accumulation of viral proteins (ICP27, ICP0, TK and VP16) was measured by immunoblotting.

## Notes

### Competing Interest Statement

The authors have declared no competing interest.

## References

1. Schoenfelder, S. and P. Fraser, Long-range enhancer-promoter contacts in gene expression control. Nat Rev Genet, 2019. 20(8): p. 437–455.

2. Marchal, C., J. Sima, and D.M. Gilbert, Control of DNA replication timing in the 3D genome. Nature Reviews Molecular Cell Biology, 2019. 20(12): p. 721–737.

3. Wang, H., M. Han, and L.S. Qi, Engineering 3D genome organization. Nat Rev Genet, 2021. 22(6): p. 343–360.

4. Yue, L., et al., Probing the spatiotemporal patterns of HBV multiplication reveals novel features of its subcellular processes. PLoS Pathog, 2021. 17(8): p. e1009838.

5. Tang, D., et al., Transcriptionally inactive hepatitis B virus episome DNA preferentially resides in the vicinity of chromosome 19 in 3D host genome upon infection. Cell Rep, 2021. 35(13): p. 109288.

6. Xiao, K., et al., RUNX1-mediated alphaherpesvirus-host trans-species chromatin interaction promotes viral transcription. Sci Adv, 2021. 7(26).

7. Yang, B., et al., 3D landscape of Hepatitis B virus interactions with human chromatins. Cell Discov, 2020. 6(1): p. 95.

8. Hensel, K.O., et al., Episomal HBV persistence within transcribed host nuclear chromatin compartments involves HBx. Epigenetics Chromatin, 2018. 11(1): p. 34.

9. James, C., et al., Herpes simplex virus: global infection prevalence and incidence estimates, 2016. Bull World Health Organ, 2020. 98(5): p. 315–329.

10. Bauer, D.W., et al., Herpes virus genome, the pressure is on. J Am Chem Soc, 2013. 135(30): p. 11216–21.

11. Brandariz-Nunez, A., et al., Pressure-driven release of viral genome into a host nucleus is a mechanism leading to herpes infection. Elife, 2019. 8.

12. McElwee, M., et al., Structure of the herpes simplex virus portal-vertex. PLoS Biol, 2018. 16(6): p. e2006191.

13. Liu, Y.T., et al., Cryo-EM structures of herpes simplex virus type 1 portal vertex and packaged genome. Nature, 2019. 570(7760): p. 257–261.

14. Brandariz-Nunez, A., S.J. Robinson, and A. Evilevitch, Pressurized DNA state inside herpes capsids-A novel antiviral target. PLoS Pathog, 2020. 16(7): p. e1008604.

15. Ojala, P.M., et al., Herpes simplex virus type 1 entry into host cells: reconstitution of capsid binding and uncoating at the nuclear pore complex in vitro. Mol Cell Biol, 2000. 20(13): p. 4922–31.

16. Newcomb, W.W., et al., Polarized DNA ejection from the herpesvirus capsid. J Mol Biol, 2009. 392(4): p. 885–94.

17. Shahin, V., et al., The genome of HSV-1 translocates through the nuclear pore as a condensed rod-like structure. J Cell Sci, 2006. 119(Pt 1): p. 23–30.

18. Roizman, B., H.D. Gu, and G. Mandel, The first 30 minutes in the life of a virus - unREST in the nucleus. Cell Cycle, 2005. 4(8): p. 1019–1021.

19. Burkham, J., D.M. Coen, and S.K. Weller, ND10 protein PML is recruited to herpes simplex virus type 1 prereplicative sites and replication compartments in the presence of viral DNA polymerase. J Virol, 1998. 72(12): p. 10100–7.

20. Knipe, D.M. and A. Cliffe, Chromatin control of herpes simplex virus lytic and latent infection. Nat Rev Microbiol, 2008. 6(3): p. 211–21.

21. Everett, R.D., et al., Formation of nuclear foci of the herpes simplex virus type 1 regulatory protein ICP4 at early times of infection: localization, dynamics, recruitment of ICP27, and evidence for the de novo induction of ND10-like complexes. J Virol, 2004. 78(4): p. 1903–17.

22. Triezenberg, S.J., R.C. Kingsbury, and S.L. McKnight, Functional dissection of VP16, the trans-activator of herpes simplex virus immediate early gene expression. Genes Dev, 1988. 2(6): p. 718–29.

23. Stern, S. and W. Herr, The herpes simplex virus trans-activator VP16 recognizes the Oct-1 homeo domain: evidence for a homeo domain recognition subdomain. Genes Dev, 1991. 5(12B): p. 2555–66.

24. Babb, R., et al., DNA recognition by the herpes simplex virus transactivator VP16: a novel DNA-binding structure. Mol Cell Biol, 2001. 21(14): p. 4700–12.

25. Douville, P., et al., Positive and negative regulation at the herpes simplex virus ICP4 and ICP0 TAATGARAT motifs. Virology, 1995. 207(1): p. 107–16.

26. Cabral, J.M., H.S. Oh, and D.M. Knipe, ATRX promotes maintenance of herpes simplex virus heterochromatin during chromatin stress. Elife, 2018. 7.

27. Hagglund, R. and B. Roizman, Role of ICP0 in the strategy of conquest of the host cell by herpes simplex virus 1. J Virol, 2004. 78(5): p. 2169–78.

28. Gu, H. and B. Roizman, The two functions of herpes simplex virus 1 ICP0, inhibition of silencing by the CoREST/REST/HDAC complex and degradation of PML, are executed in tandem. J Virol, 2009. 83(1): p. 181–7.

29. Gu, H.D. and B. Roizman, Herpes simplex virus-infected cell protein 0 blocks the silencing of viral DNA by dissociating histone deacetylases from the CoREST-REST complex. Proceedings of the National Academy of Sciences of the United States of America, 2007. 104(43): p. 1713417139.

30. Bayani, J. and J.A. Squire, Fluorescence in situ Hybridization (FISH). Curr Protoc Cell Biol, 2004. **Chapter 22**: p. Unit 22 4.

31. Xu, P. and B. Roizman, The SP100 component of ND10 enhances accumulation of PML and suppresses replication and the assembly of HSV replication compartments. Proc Natl Acad Sci U S A, 2017. 114(19): p. E3823–E3829.

32. Roizman, B., and D. M. Knipe., Herpes simplex viruses and their replication p. 2399–2459. In D. M. Knipe, P. Howley, D. E. Griffin, R. A. Lamb, M. A. Martin, B. Roizman, and S. E. Straus (ed.) Fields of Virology, 4th ed. Lippincott-Williams & Wilkins, Philadelphia, Pa. 2001.

33. Morgan, S.L., et al., Manipulation of nuclear architecture through CRISPR-mediated chromosomal looping. Nat Commun, 2017. 8: p. 15993.

34. Wang, H., et al., CRISPR-Mediated Programmable 3D Genome Positioning and Nuclear Organization. Cell, 2018. 175(5): p. 1405-1417 e14.

35. Kim, J.H., et al., LADL: light-activated dynamic looping for endogenous gene expression control. Nat Methods, 2019. 16(7): p. 633–639.

36. Dalrymple, M.A., et al., DNA sequence of the herpes simplex virus type 1 gene whose product is responsible for transcriptional activation of immediate early promoters. Nucleic Acids Res, 1985. 13(21): p. 7865–79.

37. Lu, R. and V. Misra, Potential role for luman, the cellular homologue of herpes simplex virus VP16 (alpha gene trans-inducing factor), in herpesvirus latency. J Virol, 2000. 74(2): p. 934–43.

38. Gruffat, H., R. Marchione, and E. Manet, Herpesvirus Late Gene Expression: A Viral-Specific Pre-initiation Complex Is Key. Front Microbiol, 2016. 7: p. 869.

39. Merkl, P.E., M.H. Orzalli, and D.M. Knipe, Mechanisms of Host IFI16, PML, and Daxx Protein Restriction of Herpes Simplex Virus 1 Replication. J Virol, 2018. 92(10).

40. Xu, P., S. Mallon, and B. Roizman, PML plays both inimical and beneficial roles in HSV-1 replication. Proc Natl Acad Sci U S A, 2016. 113(21): p. E3022–8.

41. Lin, J.L., et al., CRISPR-PIN: Modifying gene position in the nucleus via dCas9-mediated tethering. Synth Syst Biotechnol, 2019. 4(2): p. 73–78.

42. Shin, Y., et al., Liquid Nuclear Condensates Mechanically Sense and Restructure the Genome. Cell, 2019. 176(6): p. 1518.

43. Wang, H.F., et al., CRISPR-Mediated Programmable 3D Genome Positioning and Nuclear Organization. Cell, 2018. 175(5): p. 1405–+.

44. Kumaran, R.I., R. Thakar, and D.L. Spector, Chromatin dynamics and gene positioning. Cell, 2008. 132(6): p. 929–34.

45. Reddy, K.L., et al., Transcriptional repression mediated by repositioning of genes to the nuclear lamina. Nature, 2008. 452(7184): p. 243–7.

46. Buchwalter, A., J.M. Kaneshiro, and M.W. Hetzer, Coaching from the sidelines: the nuclear periphery in genome regulation. Nat Rev Genet, 2019. 20(1): p. 39–50.

47. Robinett, C.C., et al., In vivo localization of DNA sequences and visualization of large-scale chromatin organization using lac operator/repressor recognition. J Cell Biol, 1996. 135(6 Pt 2): p. 1685–700.

48. Finlan, L.E., et al., Recruitment to the nuclear periphery can alter expression of genes in human cells. PLoS Genet, 2008. 4(3): p. e1000039.

49. Zullo, J.M., et al., DNA sequence-dependent compartmentalization and silencing of chromatin at the nuclear lamina. Cell, 2012. 149(7): p. 1474–87.

50. Ruault, M., M. Dubarry, and A. Taddei, Re-positioning genes to the nuclear envelope in mammalian cells: impact on transcription. Trends Genet, 2008. 24(11): p. 574–81.

51. Janicki, S.M., et al., From silencing to gene expression: real-time analysis in single cells. Cell, 2004. 116(5): p. 683–98.

52. Silva, L., et al., Roles of the nuclear lamina in stable nuclear association and assembly of a herpesviral transactivator complex on viral immediate-early genes. mBio, 2012. 3(1).

53. Silva, L., et al., Role for A-type lamins in herpesviral DNA targeting and heterochromatin modulation. PLoS Pathog, 2008. 4(5): p. e1000071.

54. Everett, R.D. and J. Murray, ND10 components relocate to sites associated with herpes simplex virus type 1 nucleoprotein complexes during virus infection. J Virol, 2005. 79(8): p. 5078–89.

55. Sullivan, T., et al., Loss of A-type lamin expression compromises nuclear envelope integrity leading to muscular dystrophy. J Cell Biol, 1999. 147(5): p. 913–20.

56. Nikolova, V., et al., Defects in nuclear structure and function promote dilated cardiomyopathy in lamin A/C-deficient mice. J Clin Invest, 2004. 113(3): p. 357–69.

